# The eco-evolutionary assembly of complex communities with multiple interaction types

**DOI:** 10.1101/2025.10.29.685318

**Authors:** Gui Araujo, Miguel Lurgi

## Abstract

Identifying the mechanisms that generate structure in complex ecological communities is fundamental to understand their assembly. A comprehensive picture of how ecology and evolution act together to generate these patterns is in its infancy. We developed an eco-evolutionary model of community assembly that incorporates interaction-driven population dynamics and evolutionary processes, including speciation and the inheritance of interactions, to unveil the mechanisms of generation and maintenance of biodiversity in complex species interaction networks. Importantly, our model unpicks the effects of selection of interaction types from those of inheritance by comparing evolutionary assembly with assembly by invasion under different combinations of interaction types. We found that a cost-benefit balance of accumulating interactions separates communities into two distinct types. Interactions with weak benefits for the partners involved produce communities with sparse species interaction networks dominated by competition. Strongly beneficial interactions, on the other hand, give rise to highly mutualistic, more connected communities. Mutualism, driven by both selection and inheritance, facilitates the emergence of large yet stable communities with increased complexity. By contrasting the model results with empirical patterns from microbial communities, we identify potential drivers of the assembly of these complex ecosystems and their likely patterns of interaction. Our results provide a classification system of complex ecosystems based on their composition of ecological interactions, thus generating testable hypotheses on the conditions under which different community types (mutualistic vs. competitive) might emerge.

## Introduction

Unveiling and understanding the processes that generate and maintain biodiversity is a central focus of ecological research. Both evolutionary (e.g. speciation) and ecological (e.g. invasions) assembly mechanisms, and how they are shaped by biotic and abiotic factors such as species interactions and environmental conditions, have been shown to promote the complexity observed in natural communities [1–6]. Clarifying the relative importance of different mechanisms that foster biodiversity, while maintaining the stability of complex communities thus assembled, is fundamental to understand ecosystems [7, 8] and to inform future conservation strategies [9, 10].

Five decades ago, Robert May [11] demonstrated that ecological systems with a large number of species and high connectivity are less likely to be stable than simpler ones. This seminal finding sparked a decades-long search for the organisational features that enable natural communities to increase in complexity without compromising stability or persistence. May’s result referred to randomly connected communities (i.e., not subjected to a deterministic structure or assembly processes) represented by randomly sampled community matrices. Subsequent work has shown that particular network structures can alter the complexity-stability relationship [12], and that in empirical systems a large number of species often translates into low connectivity between them [3]. Furthermore, beyond structural patterns, the relative composition of interaction types, such as mutualism, competition, and consumer-resource interactions, has emerged as a key factor influencing the outcomes of community assembly and the stability of the resulting communities[13–17]. Only recently has theoretical research begun to explore how the interplay between different interaction types shapes the temporal assembly of complex communities, with recent studies revealing the key role of mutualistic interactions in fostering diversity (e.g. [18]). The next challenge is to uncover the mechanisms driving the emergence of different compositions of interaction types, the structural patterns of the resulting species interaction networks, and the influence of these patterns and processes on the diversity-connectivity relationships that underpin the complexity of ecological communities.

Evolutionarily, speciation is a major driver of community assembly. Speciation, acting on an existing pool of species and traits, together with subsequent selection, has been shown to recover patterns of biodiversity and network structure observed in complex model ecosystems [4, 19]. These findings suggest that speciation-driven assembly offers a key pathway for the emergence of complexity in ecological communities. Comparing speciation-driven evolutionary assembly with ecological assembly via repeated invasions helps clarify the distinct contributions of evolutionary processes to community structure [20]. Considering both speciation and invasion processes together, and their interplay, enables the investigation of two key mechanisms shaping community structure: selection of interaction types and inheritance of interactions. To date, theoretical studies of evolutionary community assembly have primarily focused on single-type species interaction networks such as food webs or mutualistic networks [4, 5, 19–25]. The combined effects of multiple interaction types have not yet been explored in evolutionary models of community assembly, despite their potential role in generating ecological complexity.

Network structure not only determines biodiversity, complexity, and stability but can also characterise distinct types of ecological communities. These structural properties often emerge from different ecological and evolutionary assembly mechanisms, as observed across diverse ecosystems [26]. For instance, in microbial communities associated to phytoplankton cells in the phycosphere, community composition is shaped by host identity and metabolic profile [27, 28]. More generally, across host-associated microbiomes, microbial succession is thought to be driven by cross-feeding interactions between microbes and with the host[29]. The discovery of universal macroecological patterns of abundance and variability in microbial communities has motivated efforts to identify general network structures that can give rise to such patterns [30, 31]. Non-interactive population dynamic models have successfully reproduced many of these patterns and have been proposed as null models for microbial community structure [30, 32, 33]. However, such models fail to capture non-random correlations in species abundances observed in these communities, a caveat that has been addressed by adding sparse and weak interactions between community members [34, 35]. Together, these findings highlight the critical role of the structure of ecological interactions in shaping microbial communities and the need to identify the assembly mechanisms that give rise to such structural organisation. Similar challenges are observed across ecosystems types and scales. For example, in mutualistic networks comprising interactions between plants and their animal pollinators, characteristic patterns of network modularity and nestedness have been suggested to arise through adaptation and niche optimisation [25] or speciation [4]. A holistic eco-evolutionary understanding of how these processes and their interplay can shed further light on ecological structure across scales.

Here, we develop a deeper mechanistic understanding of community assembly by investigating the evolutionary emergence of complex ecological networks involving multiple interaction types. We develop a dynamical model in which mutualistic, consumer-resource, and competitive interactions shape the outcomes of speciation and invasion events driving community assembly and complexity. Interactions between species determine how they influence one another, potentially triggering extinction cascades. Over time, successive assembly events and ecological filtering give rise to stable interaction network structures. We find that the strength of ecological interaction benefits defines a threshold that separates two distinct community types: one dominated by competition, and the another by mutualism. Moreover, mutualism and speciation jointly promote increasing network complexity and species diversification. Nonetheless, species richness can increase independently of mutualism and speciation if enough invasion events occur and network connecitivity is low. By analysing longitudinal data from microbial communities [30, 34], we link macroecological patterns to underlying assembly mechanisms. We hypothesise that strong benefits from ecological interactions can favour the emergence of highly interactive microbial communities that display patterns of abundance correlations characteristic of these systems. By linking eco-evolutionary assembly mechanisms to emergent community structure and observed macroecological features, our results advance understanding of the processes that shape complexity in natural communities.

## Results

We developed a step-wise community assembly model in which new species are introduced sequentially to the existing community via two distinct mechanisms: (1) evolutionary speciation and (2) invasion. We start the assembly process with small communities comprising five non-interacting species.

### Assembly by evolutionary speciation

In this scenario, new species arise as a variation of a randomly selected parent species and inherits its ecological interactions with limited fidelity. A parameter Δ defines the maximum number of interactions by which the offspring may differ from the parent species. Smaller values of Δ correspond to more constrained inheritance (i.e. greater similarity to the parent), a regime we term strong inheritance. For each speciation event, all interactions of parent species are copied into the offspring. An integer *d*, representing the total number of interaction changes, is then drawn uniformly from the interval [1, Δ]. Interactions are randomly removed and/or added from the offspring until the total number of modifications, relative to the parent, equals *d*. For example, if Δ = 5 and *d* = 3, we may remove one interaction and add two, or remove two and add one, as long as the net change amounts to *d* changes. Speciation is thus a flexible process, with the average degree of inheritance controlled by Δ: larger values lead to weaker average inheritance. If a parent species has no interactions (i.e. an isolated species), at least one interaction must be added (*d* ≥ 1) to ensure variability.

### Assembly by invasion

We then define a purely ecological scenario in which new species are introduced without any inherited interactions. Instead, each invader is assigned a connectivity value drawn uniformly from the interval [*ρ*_1_, *ρ*_2_]. This value defines the probability of forming an interaction with each existing species in the community. As invaders differ in their connectivity, community-level connectivity can shift over time due to ecological filtering / selection. We set *ρ*_1_ = 0.01 and *ρ*_2_ = 0.5 to span a range comparable to connectivity values observed in speciation scenarios.

New species are introduced once community dynamics reach equilibrium, i.e., when the dynamics governed by Eq. (1) have stabilised. Species addition can destabilise the system, driving existing species below an extinction threshold and leading to their removal from the community. The addition of a new thriving species (i.e. that does not go extinct) constitutes an assembly event. Species abundances follow ecological dynamics governed by the system of differential equations in Eq. (1), where *x*_*i*_ is the abundance of species *i* in a community of *S* species. The non-negative coefficients *p*_*ij*_, *m*_*ij*_, and *c*_*ij*_ represent the strengths of consumer-resource (+/−), mutualistic (+/+), and competitive (−/−) interactions, respectively. These coefficients determine the strength of the effect of species *j* on the density of species *i*. For consumer-resource interactions, we have 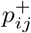 if *i* is a consumer and 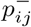 if *i* is a resource.

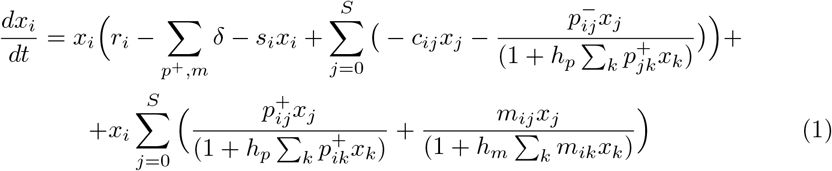

We used Type II functional responses to represent intake saturation in both mutualistic and consumer interactions. Each species *i* has a positive intrinsic growth rate *r*_*i*_ drawn from a half-normal distribution 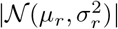. Intraspecific competition strength *s*_*i*_ is chosen such that the resulting carrying capacities *r*_*i*_*/s*_*i*_ follow a lognormal distribution. New interactions introduced via speciation or invasion are assigned a strength drawn from a half-normal distribution |𝒩 (0, *σ*^2^) |, where *σ* sets the level of interaction strength.

To capture saturation of resource intake, we assign two parameters: *h*_*p*_ for consumers and *h*_*m*_ for mutualists, representing their average handling times. Every positive interaction (whether a consumer-resource link in which species *i* is the consumer, or a mutualistic link in which *i* benefits) incurs a fixed *harvesting cost δ*. This cost is subtracted from *r*_*i*_ for each beneficial interaction, reflecting a trade-off: as a species accumulates positive links, it becomes increasingly reliant on its partners and may ultimately depend on them for survival (i.e., obligate mutualism). The balance between the interaction strength parameter *σ* and the per-link cost *δ* therefore determines the net benefit of positive interactions.

### The interplay between the strength and the cost of interactions creates a threshold that separates competitive from mutualistic communities

After 500 evolutionary assembly steps, we observed a threshold in the benefit of ecological interactions that separates two distinct community types (Fig 1). When *σ* is low and *δ* is high, resulting in weak interaction benefits, communities are dominated by competitive interactions (Fig 1C). We refer to this regime as Type 1 (T1). Conversely, when *σ* is high and *δ* is low, producing strong benefits from ecological interactions, mutualisms predominate (Fig 1A). We refer to this regime as Type 2 (T2). Consumer-resource interactions reach their highest proportion at the interface between T1 and T2, corresponding to intermediate levels of interaction benefit (Fig 1B).

**Fig. 1.**
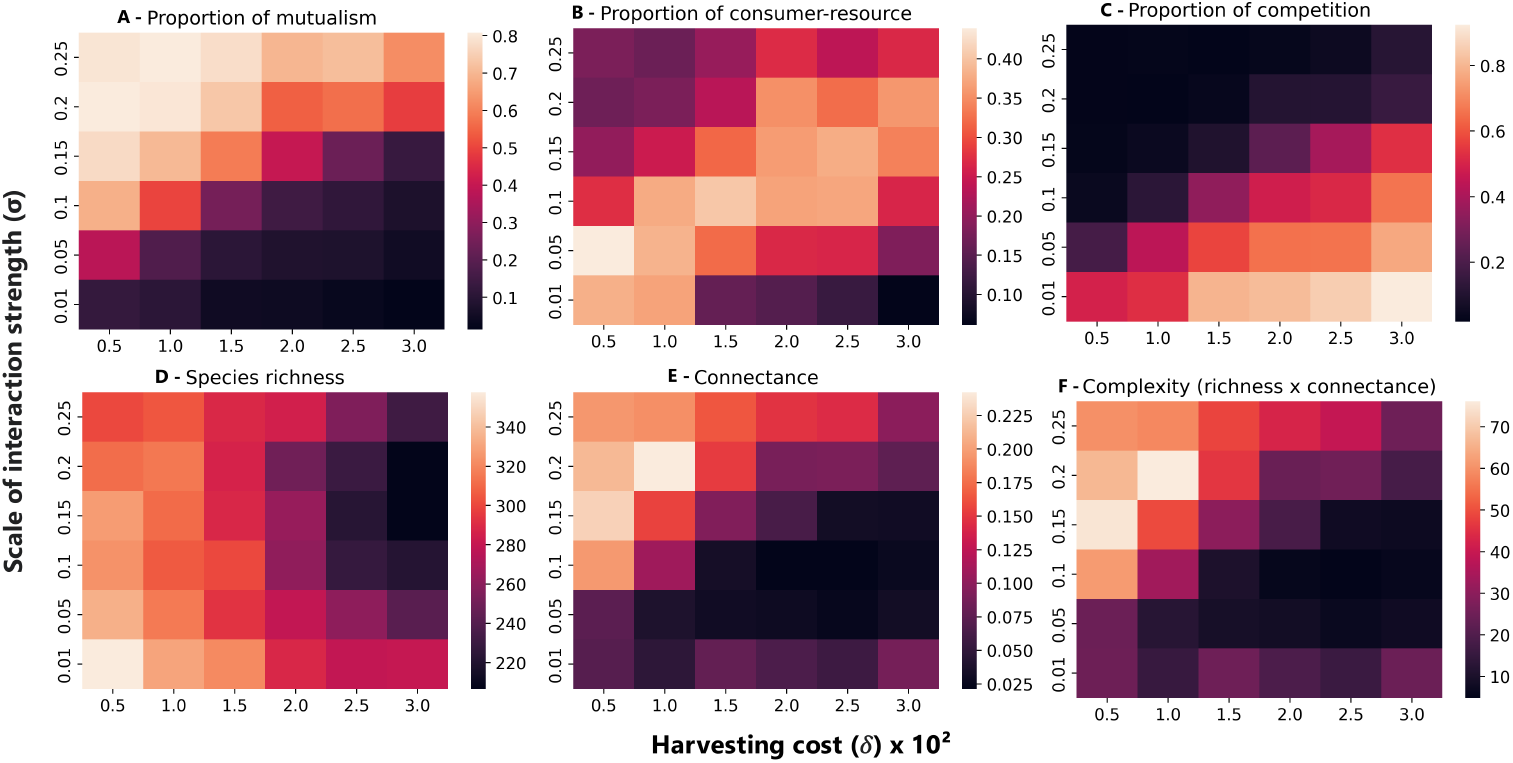
The benefit of ecological interactions creates a threshold that separates two community types. Final community structure after 500 assembly events under evolutionary assembly with strong inheritance (Δ = 5), shaped by the interplay between interaction strength (*σ*) and the cost of positive interactions or *harvesting cost* (*δ*). (A-C) The proportions of interaction types reveal a threshold separating two distinct community regimes. Type 1 communities, where interaction benefits are weak (low *σ* or high *δ*), are dominated by competition. Type 2 communities, where benefits are strong (high *σ* and low *δ*), are dominated by mutualism. Consumer-resource interactions have a higher proportion at the interface between the two regimes, particularly at low interaction costs. (D-F) Community complexity, defined as species richness multiplied by connectance, is higher in Type 2 communities and follows the same pattern as connectance, as species richness only declines under high cost and high interaction strength. Values in heatmaps represent means across 15 replicate simulations for each parameter combination. Interaction strengths were drawn from a half-normal |𝒩 (0, *σ*^2^)|. Communities started with 5 non-interacting species, and new species were introduced by speciation with Δ = 5 and initial abundances equal to the extinction threshold (10^*−*6^). 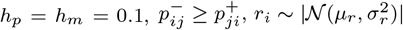.

Species richness (i.e. the number of species in the network) decreases with increasing harvesting cost *δ* (Fig 1D), while network connectance (*C* = *L/S*^2^, a measure of connectivity) reflects the threshold between community types, being higher in T2 communities (Fig 1E). As a result, community complexity, defined as the product of species richness and connectance, substantially increases in mutualism-dominated T2 communities (Fig 1F). Additionally, we found that the intensity of intraspecific competition shifts the position of the threshold (Supplementary Fig S1), indicating that the benefits of interactions must be evaluated relative to the strength of self-regulation. The existence of this threshold highlights interaction benefits as key drivers of community assembly, shaping the composition of interaction types and mediating the emergence of complexity.

### Selection and inheritance drive the composition of interaction types

Having established two distinct community types driven by interaction benefits, we next examine the role of inheritance in shaping community structure. To do so, it is necessary to distinguish inheritance effects from ecological filtering driven by the selection of interaction types that characterises the compositions of T1 and T2 communities. Both interaction-type selection and inheritance contribute to assembly under evolution, but only selection is present in assembly by invasion. By comparing evolutionary with invasion-driven assembly both considering the selection of interaction types, we can isolate the effects of inheritance.

To this end, we designed four assembly scenarios. These scenarios combine two community types (T1 and T2), two inheritance regimes (weak and strong, for evolution), and two levels of variability of interaction types (low and high, for invasion), yielding a total of eight scenarios (Fig 2A). The key distinction among these scenarios lies in how new species acquire their ecological interactions during assembly events.

**Fig. 2.**
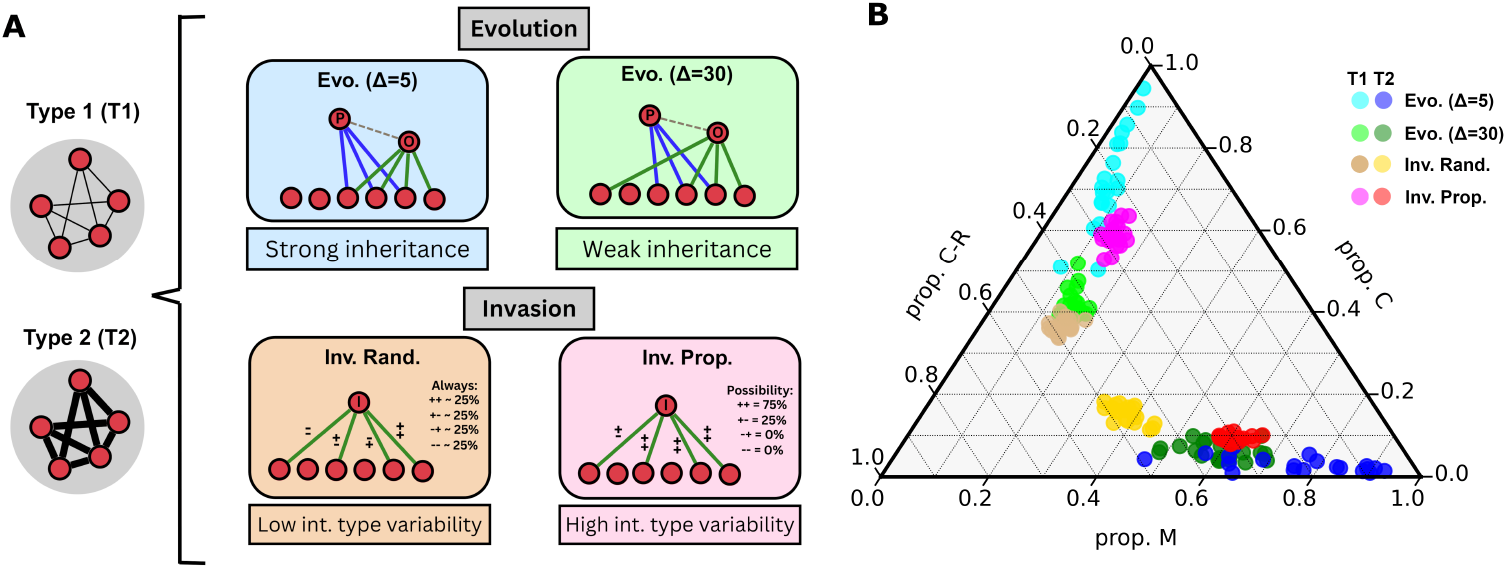
Selection and inheritance of interactions drives the composition of interaction types in assembled communities. (A) To disentangle the effects of inheritance from those of interaction-type selection, we analyse eight assembly scenarios, four for each community type (Type 1 and Type 2), spanning evolutionary and invasion processes. For evolution, we define strong (Δ = 5) and weak (Δ = 30) inheritance of interactions from *parent* to *offspring*. For invasion, we vary the level of interaction-type variability associated with the *invader* : low (random interaction types; *Inv. Rand*.) and high (random interaction-type proportions; *Inv. Prop*.). High interaction-type variability, present in both evolutionary scenarios and *Inv. Prop*., enables stronger interaction-type selection. Type 1 communities are defined by weak interaction benefits (*σ* = 0.05, *δ* = 0.025), and Type 2 communities by strong benefits (*σ* = 0.2, *δ* = 0.01). (B) Ternary plot showing the proportions of mutualistic (M), competitive (C), and consumer-resource (C-R) interactions across assembled communities. Each point represents a community at equilibrium after 500 assembly events, coloured by assembly scenario. Communities assembled under strong inheritance (Δ = 5) show more extreme separation between community types, while those assembled via *Inv. Rand*. converge toward intermediate compositions due to limited interaction-type selection. Evolutionary scenarios produce more diverse interaction-type compositions across communities. Other parameter values are the same as in Fig 1.

#### Evo. (Δ = 5)

evolution with strong inheritance. A new species arises via speciation from a randomly selected parent species. After copying the parent’s interactions, up to five additions or deletions are allowed (Δ = 5), resulting in high inheritance fidelity and close similarity between parent and offspring.

#### Evo. (Δ = 30)

As above, but with Δ = 30, offspring may differ by up to thirty interactions, producing low inheritance fidelity and greater divergence from the parent species.

#### Inv. Rand

invasion with fixed probabilities for each interaction type. Each new species is introduced with independently assigned interactions, where each link is randomly assigned to one of the four available interaction types (mutualist, competitor, consumer, or resource). Due to the random nature of this process, this will yield a probability of 0.25 for each of the interaction types. Because these probabilities are fixed, the proportion of interaction types shows low variability across invaders, limiting the scope for ecological filtering (i.e. selection cannot select species with larger proportions of specific interaction types because they do not exist). This scenario serves as a baseline and mirrors previous models of invasion-based assembly with mixed interactions [18].

#### Inv. Prop

Invasion with variable proportions of interaction types. For each new invader, the overall proportions of the four interaction types are randomly drawn from a uniform distribution. Interactions are then assigned accordingly. This introduces high variability in the species-level composition of interaction types, allowing ecological dynamics to selectively shape (via selection) the structure of community-level interactions composition.

After 500 assembly events and at ecological equilibrium, the eight scenarios produced a broad range of interaction-type compositions (Fig 2B). Across all scenarios, the threshold between T1 and T2 communities was preserved: competitive interactions dominated in T1, while mutualistic interactions predominated in T2. However, inheritance and interaction-type selection each independently amplified the divergence between community types. Strong inheritance (Δ = 5) led to the clearest predominance of the corresponding selected interaction type (consumer-resource or mutualistic), as well as the largest variability in the composition of interaction types. These results underscore the importance of species-level inheritance in shaping community-level structure and in reinforcing the effects of selection across community types.

### Evolution fosters diversity and complexity in assembled networks

To identify the drivers of community complexity, we recorded the trajectories of each assembly scenario across 500 assembly events, evaluated at ecological equilibrium (Fig 3A,C). The increase in complexity (*SxC*) was strongly dependent on community type, with T2 communities reaching substantially higher values across all scenarios. Across both community types however, both selection and inheritance independently contributed to increases in complexity. In T1 communities, *Evo. (*Δ = 30) and *Inv. Rand*. exhibited similar trajectories, due to the low connectance inhibiting selection from effectively influencing the outcome of assembly when Δ is high. By contrast, *Evo. (*Δ = 5) and *Inv. Prop*. communities reached higher average values of connectance and richness (Fig 3B), leading to a more sustained increase in complexity following a rapid initial increase (Fig 3A). Although species richness increased substantially in all communities, connectance remained consistently low. This suggests that when interaction benefits are low, resulting in competitive T1 communities, assembly tends to produce sparsely connected networks with limited complexity.

**Fig. 3.**
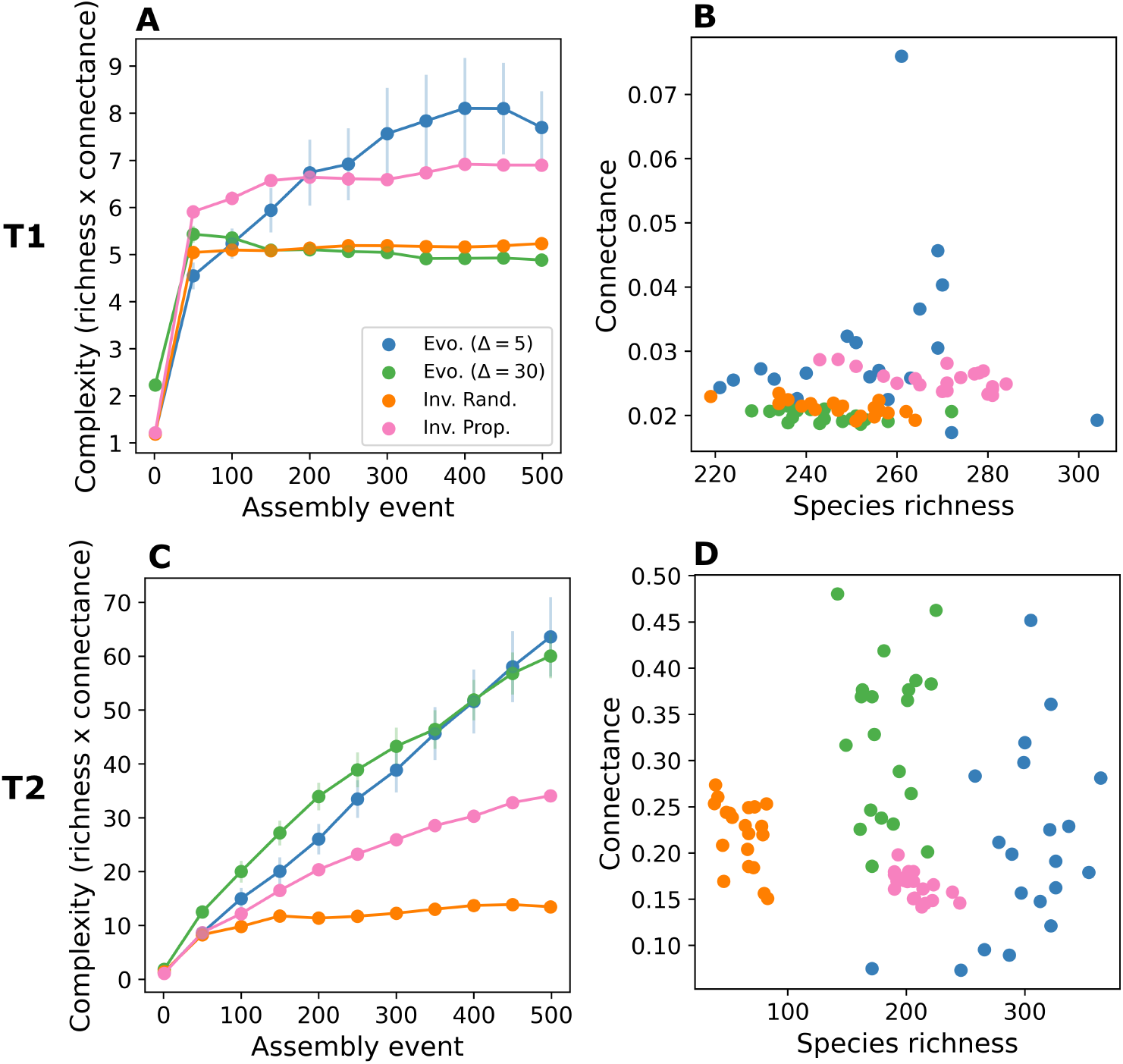
Complexity emerges from selection and inheritance of interactions. (A,C) Assembly histories showing the emergence of community complexity, defined as the product of species richness and network connectance, over 500 assembly events. Dots represent the mean of 20 replicate simulations sampled every 50 events, with vertical lines indicating the standard error 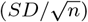. Type 2 communities consistently reach higher levels of complexity across all scenarios. In Type 1 communities, interaction-type selection increases complexity, while inheritance has minimal effect. In contrast, both selection and inheritance independently contribute to complexity in Type 2 communities. Weak and strong inheritance result in similar final complexity. (B,D) Final values of network connectance versus species richness for the communities shown in panels A and C, measured at assembly event 500. Each dot represents a single simulation. Type 1 communities attain high species richness but low connectance, whereas Type 2 communities maintain higher connectance while increasing richness. Evolutionary scenarios produce greater variability in structural outcomes. Strong and weak inheritance yield high complexity through distinct routes, either via greater connectance or greater richness. Community types were specified as in Figure 2, other parameter values are the same as in Figure 1.

In T2 communities, the *Inv. Rand*. scenario produced communities with the lowest complexity, primarily due to reduced species richness (Fig 3C,D). In contrast, *Inv. Prop*. resulted in higher complexity due to increased species richness, despite exhibiting lower average connectance (Fig 3C,D). Evolutionary scenarios had a stronger impact on assembly outcomes than invasion. Both weak and strong inheritance produced similarly high levels of complexity, the highest among all scenarios (Fig 3B), but through distinct structural pathways. *Evo. (*Δ = 5) produced communities with higher richness and lower connectance, while *Evo. (*Δ = 30) yielded the opposite pattern (Fig 3D). Thus, the strength of inheritance acts as a mechanism shaping alternative routes to complexity. Moreover, evolutionary scenarios resulted in greater heterogeneity of communities across the richness vs. connectance space, compared to invasion (Fig 3D). Overall, high interaction benefits that promote mutualism are also consistently associated with increased community complexity. Both the selection of interaction types and inheritance contributed to the predominance of mutualism and the resulting emergence of complexity.

Our results suggest that the effectiveness of inheritance in driving the emergence of complexity depends on community type, and that the degree of inheritance determines the specific pathway through which complexity arises. T2 communities exhibit more structured properties and clearer distinctions between assembly scenarios, highlighting the stronger influence of both selection and inheritance in these mutualism-dominated systems.

To further examine how network structure emerges during community assembly, we analysed the degree entropy (a measure of the evenness in the number of links across species) and network modularity in assembled communities. We quantified the relative increase in structure using a z-score metric, comparing each observed value *Z* to the average of 50 randomised communities *Z*_*r*_ with matching complexity. The score was defined as *Z*_*s*_ = (*Z* − *Z*_*r*_)*/Z*_*r*_ (see Methods for details). In T1 communities, strong inheritance (Δ = 5) produced a higher relative increase in both degree entropy and modularity. In T2 communities, both inheritance and selection contributed to increased entropy, but only inheritance led to a substantial increase in modularity (Supplementary Fig S2). These findings suggest that inheritance promotes the emergence of more modular networks with heterogeneous distributions of the number of interactions across species, both organisational hallmarks of ecological networks [36].

### Assembled communities recapitulate macroecological patterns in microbial communities

We compared our model results to empirical data from microbial communities in an attempt to identify the mechanisms, assembly processes, or community types that best approximate observed macroecological patterns in these ecosystems. We focused on patterns of relative abundance distributions found in both cross-sectional and time-series microbial community data using data from the human microbiome across different body sites (palm, gut, and mouth) and comprising 3,121 microbial types found in 1,341 samples [30]. As shown elsewhere [30, 34], these data exhibit consistent statistical patterns in the standardised log-mean abundance distribution (MAD) and in pairwise abundance correlations (Fig 4). Importantly, deviations from random expectations in pairwise abundance correlations have been linked, in part, to ecological interactions [34]. To generate data from our model outputs comparable to those from microbial communities, we ran stochastic simulations of communities assembled from equilibrium after 500 events (see Methods for details).

**Fig. 4.**
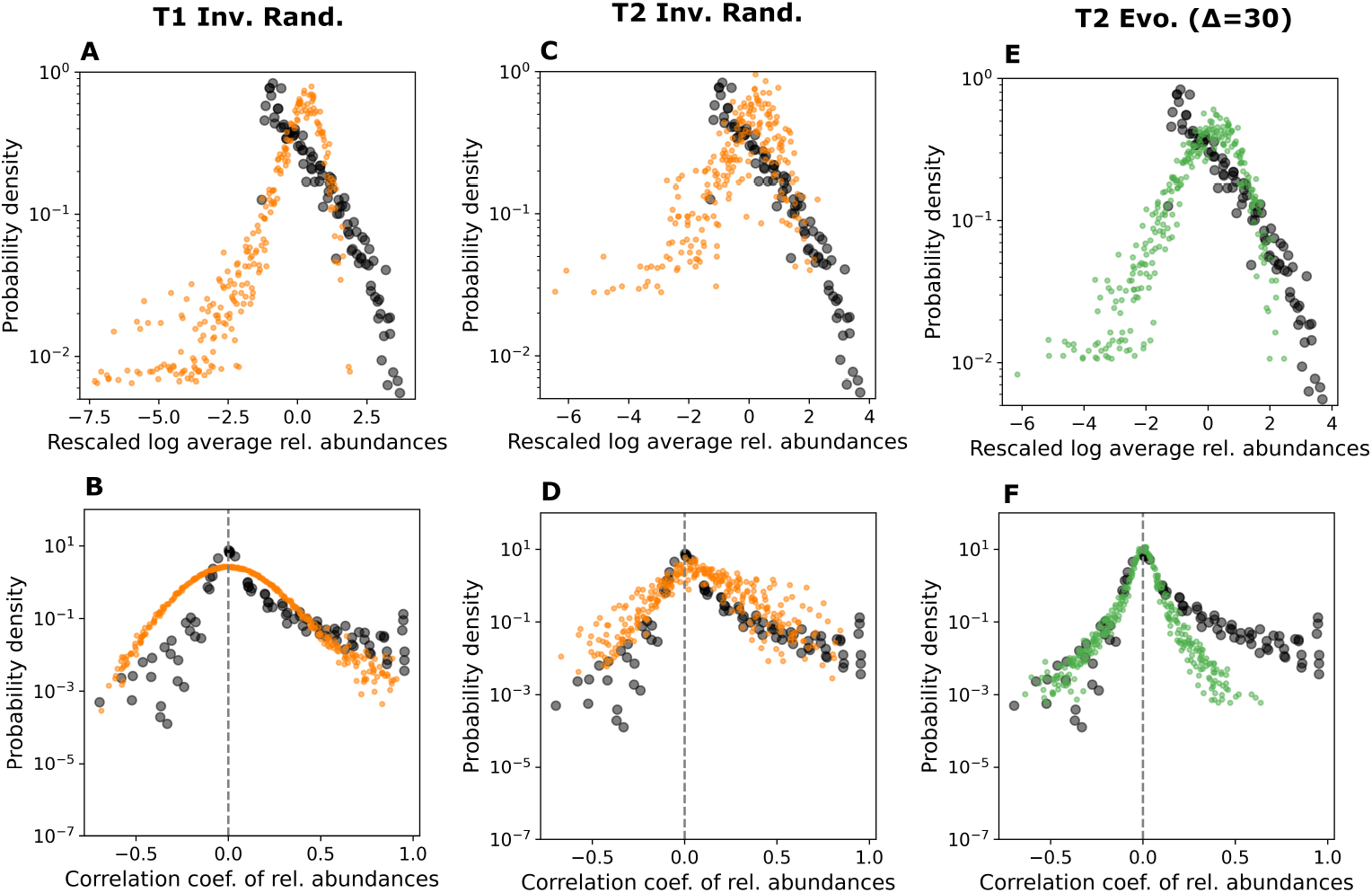
Macroecological patterns observed in microbial communities are driven by ecological and evolutionary mechanisms. We compared universal patterns from time-series data of human gut, palm, and mouth microbiomes (black dots; see Methods) to simulated communities assembled via our eco-evolutionary model. The top row (A, C, E) shows the standardised log-mean relative abundance distributions (MAD), where each point represents the probability of observing a species with a given abundance. Although real microbial abundances follow an approximately lognormal distribution, the lower tail is truncated due to sampling limitations. The bottom row (B, D, F) shows distributions of pairwise abundance correlations, where each point indicates the probability of observing a given correlation coefficient between two species as their abundances fluctuate. Coloured points represent simulation outcomes. (A-B) Type 1 *Inv. Rand*. baseline scenario. Despite the absence of interaction-type selection, the scenario produces structured MAD and correlation patterns. However, all Type 1 scenarios yield distributions nearly identical to this baseline (Supplementary Fig S3), and none approximate empirical MAD or correlation patterns. (C-D) Type 2 *Inv. Rand*. baseline. In contrast to Type 1, this scenario more closely matches the observed empirical distributions for both MAD and correlations, but fails to capture the high-abundance range in the MAD. (E-F) Type 2 weak inheritance (*Evo*. Δ = 30) scenario. This produces the best fit to empirical data, capturing more of the observable MAD and reproducing the distribution of negative correlations, although positive correlations remain underestimated. Simulations used the same parameters as in previous analyses, with 20 replicate communities sampled after 500 assembly events. Time-series fluctuations were generated by applying normally distributed environmental noise (standard deviation 0.1) proportional to species abundances, added independently to each species.

We used the *Inv. Rand*. scenario, in which interactions are assigned randomly, as a baseline for comparison. Surprisingly, all T1 scenarios produced similar relative abundance and pairwise correlation patterns (Supplementary Fig S3), as illustrated in Fig 4A,B for the *Inv. Rand*. baseline. This suggests that, in T1 communities, variation in assembly mechanisms does not translate into differences in abundance patterns, which may instead be dominated by environmental stochasticity. Moreover, none of the T1 scenarios represented well the empirical MAD or pairwise correlation patterns.

In contrast to T1, T2 communities yielded a closer resemblance to the empirical patterns observed in microbial communities. The *Inv. Rand*. baseline scenario approximates the observable range of the microbial mean abundance distribution (MAD), though it fails to capture the upper tail of highly abundant taxa (Fig 4C). It also reproduces the overall shape of pairwise abundance correlations (Fig 4D). Weak evolutionary inheritance (*Evo*. Δ = 30; Fig 4E,F) and the *Inv. Prop*. scenario (Supplementary Fig S2) yield an even closer match to empirical patterns. These scenarios better capture the higher density region of the MAD (Fig 4E) and approximate the empirical distribution of negative pairwise correlations, although they underestimate the frequency of positive correlations (Fig 4F).

Our analysis identifies T2 communities as better representations of microbial communities, highlighting strong interaction benefits as a potential mechanism underlying their macroecological abundance patterns. The presence of positive correlations in the *Inv. Rand*. baseline suggests that a more balanced mix of interaction types, as generated by random interactions, may be important for reproducing the positive pairwise correlations characteristic of microbial communities.

## Discussion

The assembly of complex ecological communities is the result of evolutionary and ecological processes acting together to generate and maintain diversity. However, our understanding of how eco-evolutionary mechanisms interact to produce such diversity remains incomplete. Here, we investigated how interaction-type selection and inheritance shape the structure of species interaction networks in a dynamic community assembly framework. We identified two distinct community regimes defined by the level of benefit species derive from ecological interactions. When interactions are weak and costly, competitive links dominate. In contrast, stronger benefits promote mutualistic communities, with consumer-resource interactions increasing in proportion at the transition between these regimes. Evolution by speciation, implementing inheritance of interactions and interaction-type selection, was found to promote community complexity. The strength of inheritance modulates how complexity emerges: strong inheritance favours increased connectance, while weak inheritance promotes species richness. These results offer insights into the mechanisms underlying natural community assembly and reveal an association between interaction benefits and macroecological patterns observed in microbial communities.

Barbier and Loreau (2019) [37] demonstrated that food webs shift from stable, self-regulated ‘pyramidal’ structures to feedback-driven ‘cascade’ dynamics depending on the balance between interaction strength and intraspecific self-regulation. Similarly, our model identifies a threshold where the balance between species self-limitation (intraspecific competition) and how much they gain from others (benefits of interactions) determines whether mainly competitive (T1) or mutualistic (T2) networks emerge. Below the threshold, strong self-regulation or high interaction costs maintain communities in a competition-dominated regime. Above it, interaction benefits become sufficiently strong to favour the emergence of mutualistic networks. These findings support a general hypothesis: community structure is governed by the interplay between interaction strength, interaction costs, and self-regulation, which together drive the emergence of distinct and dynamically stable configurations.

The ability to manage energy and resources to sustain interactions has been proposed as a key driver of ecological diversity [38]. This supports the hypothesis that variation in the cost-benefit balance of interactions gives rise to distinct community types. Mooij et al. (2024) [39] showed that Lotka-Volterra competitive communities with sparse, weak interactions can stably support unlimited species richness. These findings align with our results for Type 1 communities, which we propose emerge through assembly processes that favour competition over other interaction types. Previous studies have shown that both the mix of interaction types and their cost-benefit characteristics influence community stability and structure [14, 16]. Our results demonstrate how these characteristics can be set to emerge ecological complexity, with high interaction benefits and the selection of mutualism appearing as important mechanisms. When speciation is included, these mechanisms lead to even higher complexity in Type 2 communities. These findings highlight the joint contribution of evolution and the structure of ecological interactions as drivers of community complexity. Surprisingly, high complexity emerged alongside stronger interactions, challenging May’s classical view of random communities [11, 12] and demonstrating that highly connected, strongly interactive systems can remain stable. Although May’s inequality is true, we argue that a randomly defined community matrix (i.e. the Jacobian) do not reflect complex communities with saturating positive interactions. Moreover, our results suggest that inheritance is a key driver of modularity, which is a property associated with stability in mutualistic and microbial communities [25, 40, 41].

In addition to promoting complexity and modularity, inheritance was essential for generating diverse community outcomes in interaction composition, connectance, and species richness. This variability highlights evolutionary processes as key drivers of local and regional diversity, generating novelty in the co-occurrence patterns among communities. This conclusion aligns with the geographic mosaic theory of coevolution, which posits that local evolutionary dynamics generate population-level heterogeneity [1]. Our results show that local speciation, through species interactions alone, can drive divergent community outcomes, even in the absence of abiotic or trait-based heterogeneity. This further supports the role of priority effects and historical contingency in the evolutionary assembly of microbial communities [42, 43]. Specifically, we find that the degree of inheritance shapes distinct pathways to complexity, favouring either species richness or network connectance. This richness-connectance trade-off has been observed in microbial systems [44], and we propose that inheritance strength acts as an evolutionary mechanism driving this pattern.

Differences between the *Inv. Rand*. and *Inv. Prop*. scenarios underscore the key role of interaction-type selection in shaping community assembly. However, successful invasions require an external supply of interaction-type diversity or a precise composition matching community demands, which may be rare or unreliable conditions. We argue that interaction-type selection is more characteristic of evolutionary assembly. Unlike invasion, evolution more reliably generates variability internally, through diversification and inheritance. Thus, even ecological filtering of interaction types may be primarily driven by evolutionary processes. Consequently, the influence of inheritance through speciation may be greater than suggested by the differences between the *Inv. Prop*. and speciation scenarios.

Camacho-Mateu et. al. (2024) [34] found that sparse and weak interactions closely reproduce macroecological patterns observed in microbial communities. However, those configurations emerged from parameter optimisation rather than from a generative process based on internal community dynamics. Instead, we investigated the macroecological patterns exhibited by communities assembled via eco-evolutionary processes. While our results do not fully replicate empirical patterns, they reveal additional pathways through which these patterns may be approximated. Surprisingly, our results suggest that, when considering community assembly, strong and abundant interactions characterise microbial communities exhibiting such patterns. Moreover, previous studies linked phylogenetic distance with positive microbial abundance correlations [45]. We found that weak inheritance in T2 communities approximates empirical patterns well, except for positive correlations. This, combined with previous findings, suggests that modifying the implementation of weak inheritance, e.g. by altering how interactions evolve or incorporating taxon-specific traits, could improve alignment with empirical data. We also found that introducing species with randomly defined interactions can generate strong positive correlations, offering an alternative explanation for observed pairwise patterns in microbial communities. Future work may build on these results to explore additional mechanisms that help explain the remaining variability in microbial macroecological patterns.

We highlight the contrast between inheritance through speciation and ecological filtering through selection of interaction types as distinct drivers of eco-evolutionary community assembly. Previous models incorporating mixed interaction types have examined invasion-driven assembly, similar to our *Inv. Rand*. scenario, and identified mutualism as a key factor in community persistence [18]. We extend these findings by isolating the contribution of interaction inheritance to community assembly. This distinction is an important step toward disentangling the relative contributions of ecological and evolutionary processes in community assembly. Disentangling these contributions is especially relevant for microbiomes, which are complex communities assembled in close phylogenetic and life-long association with host individuals [26, 46]. These systems often reflect long-term coevolution among microbes and between microbes and their hosts [47], making evolutionary processes central to understanding microbiome structure and function. Future work could build on our framework to explore the mechanistic assembly of both free-living microbial communities and host-associated microbiomes to further characterise the action of evolutionary mechanisms. In this context, considering host communities alongside their associated microbes within a metacommunity framework, as recently proposed [26, 48, 49], may offer deeper insight into the mechanisms underlying the eco-evolutionary assembly of complex coevolved symbioses, highlighting the importance of dispersal and transmission modes.

Our results provide insights into the processes driving the emergence of complexity during eco-evolutionary community assembly. Key mechanisms include the strength of ecological benefits, the degree of inheritance, and the selection of interaction types, which determine whether competition or mutualism predominates. Understanding both the network structures and underlying mechanisms supporting stable, complex, and diverse communities is essential to explain their persistence in nature. This knowledge is also critical for predicting their responses to environmental change. In this way, our work advances ecological theory and supports efforts to maintain and restore complex ecological systems.

## Methods

### Simulations

All simulations begin with five species with no initial interactions. Species abundances at initialization are uniformly drawn from the interval [0, 0.02]. A species is considered extinct and removed from the system if its abundance falls below a threshold of *x*_ext_ = 10^*−*6^. Each simulation proceeds for 500 assembly events. Assembly occurs only after the system reaches ecological equilibrium, defined as a relative change in all species abundances of less than 0.01% over ten time steps of size 0.01. Intrinsic growth rates *r*_*i*_ are sampled from a normal distribution 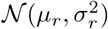 with *µ*_*r*_ = *σ*_*r*_ = 0.1. Intraspecific competition coefficients *s*_*i*_ are determined such that carrying capacities *K*_*i*_ = *r*_*i*_*/s*_*i*_ follow a lognormal distribution with underlying normal parameters *µ* = −2.2, *σ* = 0.5; specifically, *s*_*i*_ ∝ *r*_*i*_*/* log𝒩(−2.2, 0.5^2^). In evolutionary scenarios, *r*_*i*_ and *s*_*i*_ are not inherited, reflecting their dependence on species-specific environmental conditions and focusing inheritance on interaction structure only. Simulations use Type II functional response on positive interactions with constant handling times set as *h*_*m*_ = *h*_*p*_ = 0.1. When new interactions are introduced, their strengths are drawn from a half-normal distribution: |𝒩(0, *σ*^2^)|, where *σ* controls the typical strength of interactions. In evolution scenarios, inherited interaction strengths are subject to small stochastic variation, given by *w*^*′*^ = *w* + (0, [0.05*w*]^2^), where *w* is the parent’s strength and *w*^*′*^ is the offspring’s. For consumer-resource interactions, when the benefit to the consumer exceeds the cost to the resource due to random sampling (i.e. when 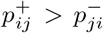), we enforce symmetry by setting 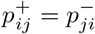. After the first 20 assembly events, all isolated species (i.e. those with no interactions) are removed after each subsequent event. Numerical integration of ecological dynamics (Eq 1) is performed using the *odeint* function from the SciPy library in Python [50].

### Analysis

For each combination of *δ* = ({0.005, 0.01, 0.015, 0.02, 0.025, 0.03} and *σ* = {0.01, 0.05, 0.1, 0.15, 0.2, 0.25}) in the heatmaps, we ran 15 independent simulations using the strong inheritance evolution model (Δ = 5), each consisting of 500 assembly events. Final community properties were averaged across these replicates and used to generate the heatmap data. To analyse the temporal dynamics of community composition (Fig 3), we ran 20 replicate simulations per scenario. Variables were recorded every 50 assembly events, and both the mean and standard error (defined as the standard deviation divided by the square root of the number of replicates) were computed and shown in the plots. Except for the heatmaps, all other results were based on these 20 replicates per scenario. Type 1 and Type 2 communities were defined by parameter combinations (*δ* = 0.025, *σ* = 0.05) and (*δ* = 0.01, *σ* = 0.2), respectively (which are contained in the heatmaps).

### Network metrics

We evaluated two structural properties of the assembled ecological networks: degree entropy and modularity. *Degree entropy:* quantifies the heterogeneity of the network’s degree distribution *P* (*k*), where *k* denotes the number of interactions (degree) of a node (species). We computed the entropy using the unweighted network (i.e., considering only presenceabsence of links), as *H* = −Σ_*k*_ *P* (*k*)*ln*(*P* (*k*)). *Modularity:* is a standard network metric that quantifies the extent to which nodes form communities, i.e., groups of species that are more densely connected internally than to other parts of the network [51]. We calculated modularity from the unweighted interaction network using 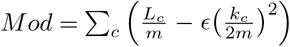 where *c* indexes each community (or module), *L*_*c*_ is the number of intra-community links, *k*_*c*_ is the sum of degrees of nodes in community *c*, and *m* is the total number of edges in the network. The resolution parameter *ϵ* was set to 1. All network analyses were performed using the Python package **NetworkX** [52], and community detection was carried out using the *Louvain algorithm* [53]. *Effective increase:* To facilitate comparison across scenarios, network metrics (degree entropy and modularity) were expressed as their effective increase relative to corresponding random networks. Specifically, for a given metric *Z*, we computed its effective increase *Z*_*s*_ as 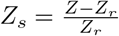, where *Z* is the observed value in the assembled network, and *Z*_*r*_ is the mean value obtained from 50 Erdős-Rényi random networks with the same number of species *S* and connectance *C*. This measure represents the percentage increase of the observed network property relative to what would be expected by chance.

## Data analysis

We utilised publicly available longitudinal human microbiome data from the EBI Metagenomics portal (project ERP021896) [54, 55], which includes samples from the tongue, palms, and gut. This is the same longitudinal dataset used by Grilli (2020) [30] to investigate macroecological patterns. We retained only samples containing at least 10,000 sequencing reads and computed species’ relative abundances accordingly. To generate comparable simulation data, we reused the same replicate simulations from Figs 2 and 3, starting from the final equilibrium state reached after 500 assembly events. We modelled species abundance fluctuations using stochastic differential equations with environmental noise. Specifically, we added independent Gaussian noise to each species, scaled proportionally to its abundance, with a standard deviation of

Integration was performed using the Euler-Maruyama method, and abundances were recorded at evenly spaced, non-overlapping time intervals.

## Software availability

Computer code developed to implement the model and execute the numerical simulations used to produce the results presented in this study is available on Zenodo: 10.5281/zenodo.15621028 and on GitHub: github/eco-evo-network-assembly.

## Supplementary information

Supplementary material is available online and includes additional figures referenced in the main text.

## Acknowledgements

This project was supported by the Leverhulme Trust through Research Project Grant # RPG-2022-114 to ML.

## Competing interests

The authors declare no competing interests.

## Author contributions

ML and GA conceived and designed the study. GA implemented the model and performed numerical simulations and analysis. ML and GA wrote the manuscript.

## Notes

https://zenodo.org/records/15621028

### Competing Interest Statement

The authors have declared no competing interest.

