## Supplementary Material for "The eco-evolutionary assembly of complex communities with multiple interaction types"

### Supplementary material for the paper '*The eco-evolutionary assembly of complex communities with multiple interaction types*'

Gui Araujo[1] & Miguel Lurgi[1,*]

[1]Department of Biosciences, Swansea University, SA2 8PP, UK.

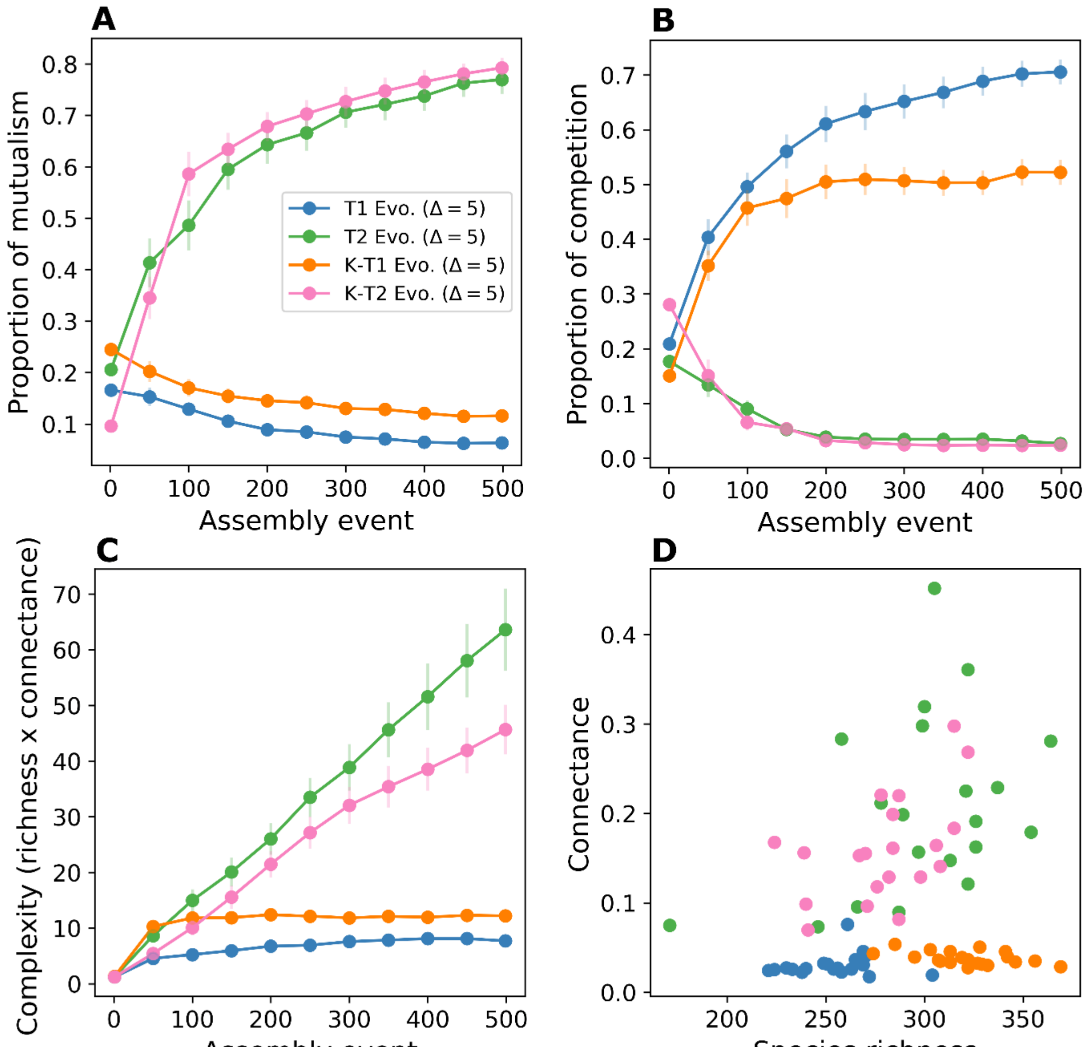


Figure S1. **The scale of intraspecific competition establishes the threshold of benefits of interactions.** The plots show the average values of 20 samples (i.e. replicated simulations of the model) for every 50 assembly events (dots) and the vertical lines show the standard errors ($SD/\sqrt{n}$). We simulate the effect on community types of changing the mean of intraspecific competition, which rescales the benefits of interactions (i.e. interaction strength σ in relation to the cost of positive interactions δ) for assembly by evolution ($\Delta = 5$). T1 and T2 curves (blue and green) are Type 1 and 2 communities, respectively defined by ($\sigma = 0.05, \delta = 0.025$) and ($\sigma = 0.2, \delta = 0.01$), as in Figure 2. The K-T1 curve is originally defined as a T2, but it also has a greater average intraspecific competition, with $s_i \propto r_i/[log\mathcal{N}(-4, 0.5^2)]$ instead of $s_i \propto r_i/[log\mathcal{N}(-2.2, 0.5^2)]$. Increasing the intraspecific competition from T2 to K-T1 transforms the outcome into a type 1 community (green and orange curves). The same applies to T1 and K-T2, in which decreasing the intraspecific competition to $s_i \propto r_i/[log\mathcal{N}(-0.4{,}0.5^2)]$ transforms the outcome into a Type 2 community (blue and magenta curves). (A-B) The characteristic outcome of interaction-type composition of both community types flips by changing $s_i$. (C-D) Complexity and its components, richness and connectance, also flip accordingly. Parameter values are the same as in Figure 2.

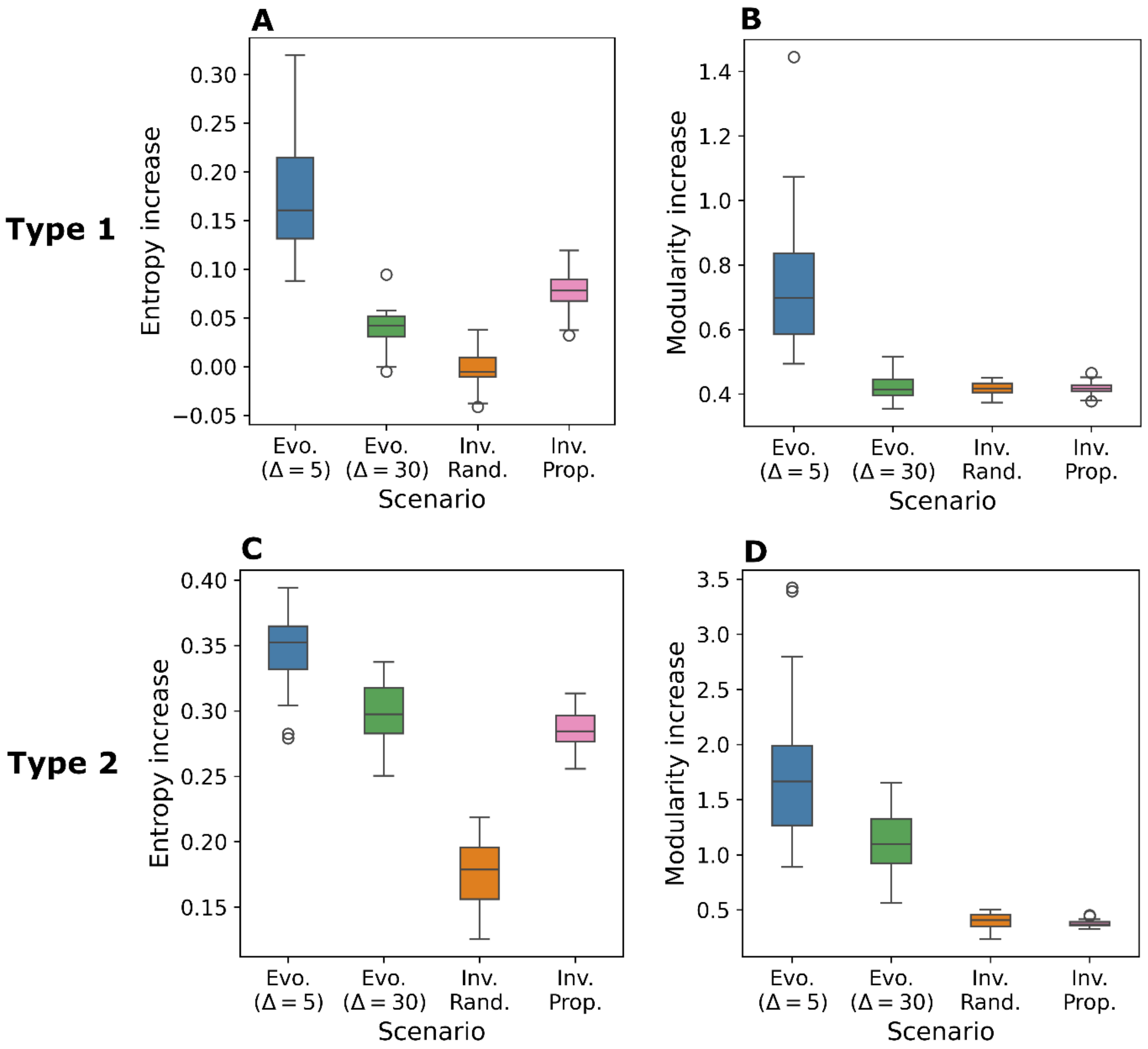


Figure S2. **Network metrics.** Degree entropy and modularity of 20 simulation samples after 500 assembly events, for each scenario (details in Methods). Metrics were analysed as the relative increase from the average random network with the same complexity, calculated with 50 samples. Values were subtracted by the random average and then divided by it, resulting in a relative increase and measuring how much is driven by the structure generated in the assembly process. (A-B) In Type 1 communities, strong inheritance is the main factor driving the increase of both entropy and modularity. (C-D) In Type 2 communities, the same holds for modularity, with weak inheritance being relevant as well, but ecological selection is also a substantial driver of entropy. Parameter values are the same as in Figures 1 and 2.

#### **Complementary data analysis scenarios (Figs S3 and S4)**

We used empirical data from the human microbiome to evaluate how well each model scenario replicates macroecological patterns observed in microbial communities. These patterns are considered universal and are illustrated here using time-series data from gut, palm, and mouth microbiomes across multiple individuals (black dots). In all panels, the top plots show the distribution of standardised log-mean relative abundances (mean abundance distributions, MAD), with each dot representing the probability of observing a species with a given abundance. Empirical MADs approximate a lognormal distribution (continuous line), although the lower tail is truncated due to detection limits in sampling. The bottom plots show the distribution of pairwise correlation coefficients between species' relative abundances, where each dot indicates the probability of observing a given correlation as abundances fluctuate over time. Coloured dots represent results from simulations. Each figure corresponds to a different community type, displaying scenarios not shown in Fig 4. Simulations were performed using the same parameter values as in the main analyses. We generated 20 replicate communities per scenario, each assembled through 500 events. To produce time-series reflecting stochastic fluctuations around equilibrium, we added normally distributed environmental noise (standard deviation 0.1) independently to each species, scaled to its abundance.

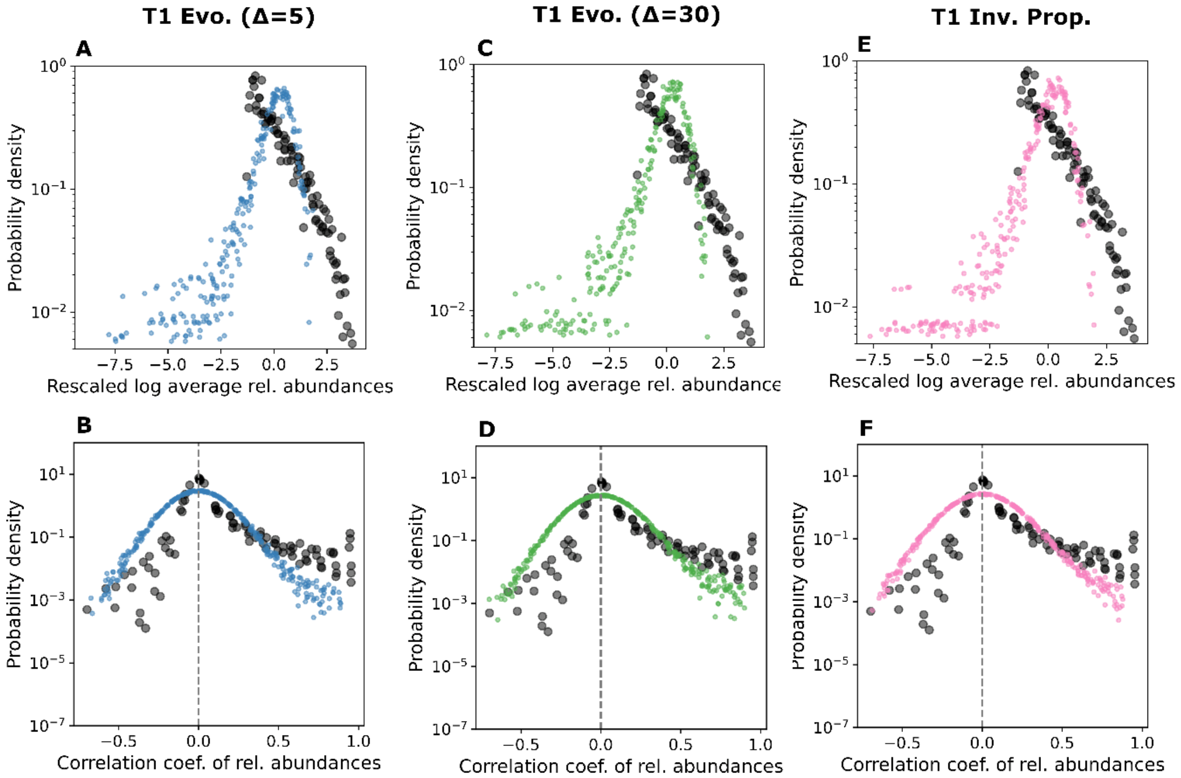

Figure S3. **Macroecological patterns for other scenarios of Type 1 communities.** (A-B) Strong inheritance. (C-D) Weak inheritance. (E-F) Invasion with variable interaction-type proportions.

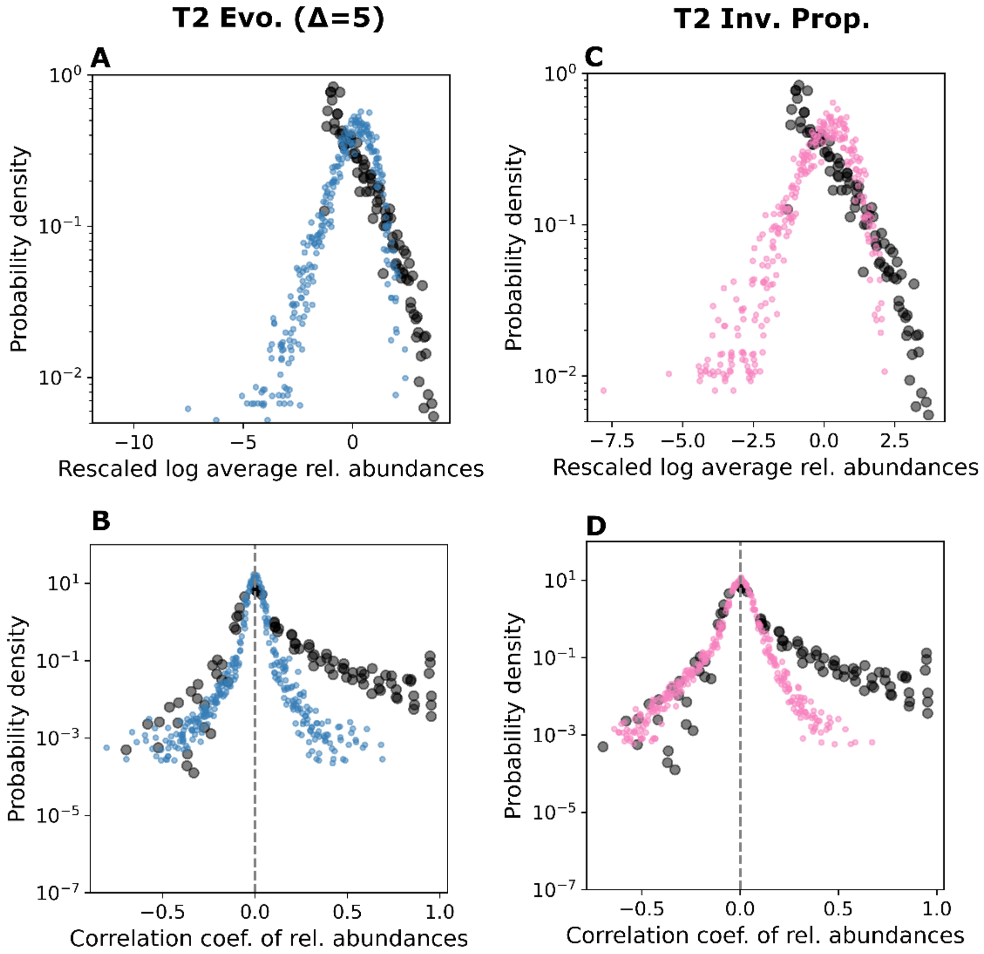


Figure S4. **Macroecological patterns for other scenarios of Type 2 communities.** (A-B) Strong inheritance. (C-D) Invasion with variable interaction-type proportions.
